# Predictive remapping and allocentric coding as consequences of energy efficiency in recurrent neural network models of active vision

**DOI:** 10.1101/2025.05.22.655555

**Authors:** Thomas Nortmann, Philip Sulewski, Tim C. Kietzmann

## Abstract

Despite moving our eyes from one location to another, our perception of the world is stable - an aspect thought to rely on predictive computations that use efference copies to predict the upcoming foveal input. Are these complex computations genetically hard-coded, or can they emerge from simpler principles? Here we consider the organism’s limited energy budget as a potential origin. We expose a recurrent neural network to sequences of fixation patches and saccadic efference copies, training the model to minimise energy consumption (preactivation). We show that targeted inhibitory predictive remapping emerges from this energy efficiency optimization alone. As furthermore demonstrated, this computation relies on the model’s learned ability to re-code egocentric eye-coordinates into an allocentric (image-centric) reference frame. Together, our findings suggest that both allocentric coding and predictive remapping can emerge from energy efficiency constraints during active vision, demonstrating how complex neural computations can arise from simple physical principles.

## Introduction

Although our eyes shift fixation locations approximately three times per second, we perceive the world as stable and continuous — a surprising phenomenon given the substantial changes in visual input that occur across saccadic eye movements. This “hard binding problem” (Cavanagh et al., 2010) represents a fundamental challenge in vision science. To address it, predictions of foveal input from upcoming eyemovements are thought to play a crucial role through predictive remapping (Golomb & Mazer, 2021). These predictions rely on efference copies encoding planned eye movements to anticipate the visual consequences of saccadic eye movements. Behavioural evidence shows that this mechanism enables optimal transmission and integration of visual features across saccades (Fabius et al., 2020; Ganmor et al., 2015; Harrison et al., 2013; D. He et al., 2017; Wolf & Schü tz, 2015). A broad network of brain areas is involved in this process (Duhamel et al., 1992; Knapen et al., 2016; Nakamura & Colby, 2002; Neupane et al., 2016; Umeno & Goldberg, 1997; Walker et al., 1995), with various proposed mechanisms for remapping peripheral information to foveal locations (Arkesteijn et al., 2019; Cavanagh et al., 2010; Golomb & Mazer, 2021; Melcher, 2007; Neupane et al., 2016, 2020; Rolfs, 2015). While the exact mechanisms of predictive remapping are still a topic of debate, what remains less controversial is the presence of (pre-)saccadic predictive computations that facilitate a seamless visual handover.

The central importance of stable perception despite eyemovements and the complexity of the computations involved raise intriguing questions about their origins: are these processes and underlying connectivity motifs genetically hardcoded, or could they emerge from simpler principles? Energy efficiency stands as a fundamental constraint in neural computation and a cornerstone principle in computational neuroscience (Lennie, 2003; Sterling & Laughlin, 2015). This principle manifests across multiple neural phenomena, from sparse coding—where information is represented by few active neurons (Olshausen & Field, 1996)—to predictive coding, where predictable sensory inputs are actively inhibited (Rao & Ballard, 1999; Friston, 2005). Our study explores whether the sophisticated neural mechanisms that maintain perceptual stability across saccades might similarly emerge from these basic energy constraints without requiring specialized genetic coding.

Previous work has demonstrated that predictive visual computations can arise in systems optimised for energy efficiency (Ali et al., 2022). However, while Ali et al. (2022) studied temporally predictable digit image sequences, the mechanism underlying predictive remapping across saccades is decisively more complex: relative spatial information about saccadic targets needs to be used to flexibly transfer information across space, e.g. from periphery to fovea. Moreover, our approach extends this previous work by utilizing naturalistic stimuli and mirrors the spatial stochasticity of human-like fixation behavior, providing a more ecologically valid test of whether energy efficiency can drive complex spatial computations. Hence it remains unclear whether the intricate task of predictive remapping can similarly be found as an emergent phenomenon driven by energy efficiency, without the need for intricate architectural design.

Here, we approach this question by developing a model framework in which a recurrent neural network (RNN) is subjected to sequences of fixation patches from natural scenes alongside their corresponding (relative) saccadic efferent copies (Fig 1). The RNN’s objective is to optimise its synaptic weights to minimise firing rates and synaptic transmission, approximating a substantial part of the energy costs related to neural information processing. This energy efficiency objective is implemented by minimizing the mean absolute preactivation (the sum of inputs to a model unit before the activation function is applied) across all units and time steps, while we maintain excitatory input to the network. In combination this aims to prevent costly overinhibiton or a learned network-shutdown in favour of energy-efficiency.

**Figure 1:**
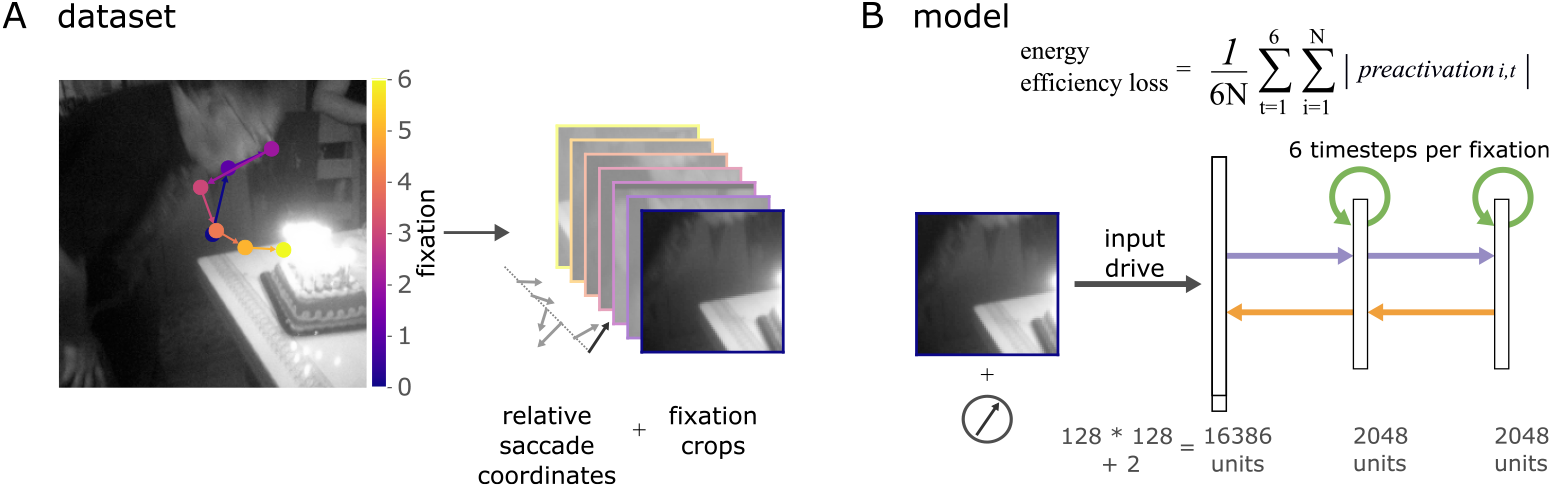
Minimizing preactivation in response to human-like saccade sequences. **(A)** Generation of fixation crop sequences with relative saccadic coordinates from the MS-COCO dataset (Lin et al., 2015), sequences based on DeepGaze III (Kü mmerer et al., 2022). **(B)** Schematic of the RNN architecture alongside depiction of the energy-efficiency loss function. The model receives excitatory input drive directly connected to the first layer (fixed, no learnable parameters) while computing through hidden layers with bottom-up (purple), lateral (green), and top-down (yellow) connections. Energy efficiency is modeled by minimizing mean absolute unit preactivation computed over all units (N) and all six timesteps, which encourages sparse neural activity without excessive inhibition. The fixed excitatory input prevents the network from learning to ignore inputs. This combination aims to support the emergence of targeted inhibitory predictions rather than costly over-inhibition or network shutdown.

To elucidate whether minimising a network’s energy expenditure, in context of saccadic eye-movements, can yield signatures of predictive remapping, we (i) compared the energy efficiency loss of model predictions against several control conditions, such as predicting the average training set luminance, the average training set fixation crop and a static repetition of the previous crop as predictor, and (ii) visualise and analyse the model-internal drive, demonstrating targeted inhibition. Finally, by performing diagnostic read-outs and in-silico lesion studies, we further reveal the network’s learned strategy to implement such predictive remapping. We find that the network learned to map the sequence of relative saccade targets into an allocentric reference frame and, importantly, that the allocentric-coding units are driving the targeted prediction.

## Results

### Inhibitory predictive remapping emerges as a consequence of energy efficiency

We trained a recurrent neural network (RNN) architecture with two hidden layers to minimise its energy consumption, implemented as minimising unit preactivation (Ali et al., 2022) while being presented with excitatory sensory input (sequences of fixation crops) and the corresponding relative saccade coordinates. Fixation sequences were generated on natural scenes sourced from the MS-COCO dataset (Lin et al., 2015; see Fig 1A) using Deepgaze III predictions of human fixation behaviour (Kü mmerer et al., 2022). Each sequence consisted of seven fixation crops, and each crop was processed for six model time steps (Fig 1B). In our analyses, we primarily focus on the computations/loss values recorded at the first model time step of each fixation crop. This timepoint enables us to investigate the model’s behaviour while transitioning from one fixation location to the next.

To assess the performance of our model, we compared its energy consumption against a series of controls. First, we contrasted the energy-efficiency trained RNN against static predictions derived from the training set (see Fig 2A). In one variant, the control model was set to subtract the average luminance of all images in the training set. We find the loss of the energy-optimised RNN to be significantly lower (*p <* .001, see Fig 2B). The second static control used the average crop, computed over all crops in the training set, as inhibitory prediction. Again, our RNN model loss was significantly smaller (p *<* .001, see Fig 2B). An additional non-static control to probe spatial biases in the dataset used the crop at the current fixation location of the average image in the training set, which controls for spatial dataset-biases (Fig 2A). Our RNN model loss was significantly lower than this control condition (p *<* .001, see Fig 2B). These controls demonstrate that the model outperforms non-informative predictions and is sensitive to scene-specific characteristics.

**Figure 2:**
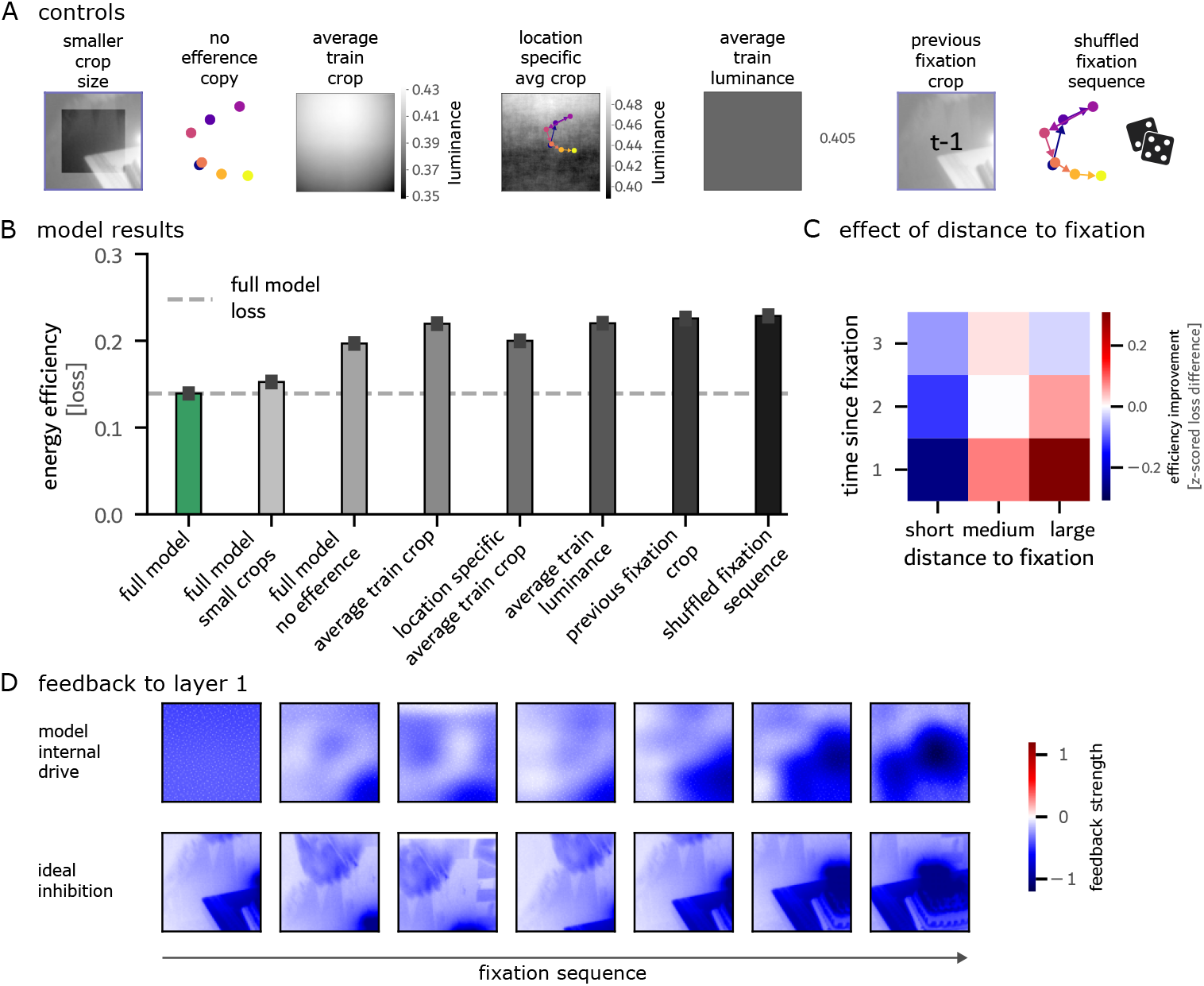
Inhibitory predictive remapping as a consequence of energy efficiency. **(A)** Illustration of the seven control conditions used for performance comparison: training with smaller crops (56 % smaller), training without efference copies, location specific average crop, average crop, average luminance value, previous fixation crop and testing with shuffled fixation sequences **(B)** Evaluation of energy efficiency loss in the RNN model and control conditions, highlighting the model’s predictive capabilities. **(C)** Mean normalised loss (z-scored by average test set loss) plotted as a function of two variables: the spatial distance from previous fixation locations and the temporal distance (number of fixation steps) since the location was last visited. **(D)** Example of the trained RNN’s internal feedback to layer 1 (upper row), together with the ideal inhibition defined as the inversion of the fixation crop (bottom row). Optimised for energy efficiency, targeted inhibition emerges in the trained RNN model. While smooth, the inhibitory patterns align with the ideal inhibition.

Next, we asked to what extent the fixation dynamics on the scene informed the energy efficiency-based predictions. To this end, we contrasted our model’s predictions against two scene-specific control conditions (Fig 2A). First, we tested against a model that performs inhibitions based on a repetition of the last fixation crop, which are then used as predictions for the upcoming crop. Our RNN model loss was lower than this previous fixation control (*p <* .001, see Fig 2B), indicating the RNN learned to adapt its predictions in response to upcoming saccadic targets on a given scene. We then ran a fifth control in which we produced a mismatch between the efference copies given to the model and the corresponding change in visual information that results from the saccade. To do so, the sequences of fixation coordinates were shuffled, while maintaining the original fixation crop sequence. Subsequent to this coordinate shuffling, we recomputed the relative saccade positions to be fed to the model. Our RNN model loss that relied on the original sequence of efference copies was lower than in the control condition with shuffled position-crop pairs (*p <* .001, see Fig 2B). This suggests that the RNN successfully couples efference copy information to visual information at saccade targets a strong indication that it performs inhibitory predictive remapping. To probe whether our model architecture could learn similar predictions in the absence of spatial information, we trained the same model without concatenating the efference copies to the input (Fig 2A). Our RNN model loss was lower than in this control model (*p <* .001, Fig 2B). This indicates the models need to rely on efference copies to achieve good spatial predictions. To investigate the model’s reliance on the overlap of fixation crops, we implemented a control model with the same architecture except a visual field reduced by 56% (Fig 2A) to reduce overlap between adjacent fixations. Despite this control model having a larger energy efficiency loss than our model, it outperformed all controls (all *p*s *<* .001, Fig 2B), indicating that the learned predictions do not solely rely on fixation overlap.

Probing the spatial memory abilities of our model, we analysed whether the energy efficiency loss changes for fixations based on if a spatially close fixation was fixated earlier. The mean loss decreases in cases where one of the last two fixations was close to the current fixation and increases if previous fixations were far away (Fig 2C). As the effect is true not only for the last but also second to last fixation, a limited form of spatial memory by the RNN is suggested.

In analysing the model more closely, we observe that, to achieve energy-efficiency in face of excitatory input, the overall top-down feedback to layer 1 of our trained RNN was inhibitory (feedback over test set crops: *µ* = −0.39, 99% CI = [−0.78, −0.09], n = 86142). To better understand the spatial specificity of this model behaviour, we next visualised the model-internal drive across sequences of eye-movements. Visual inspection of these inhibitory signals suggests that topdown feedback is spatially specific to the saccadic target locations, rather than being a global, spatially unspecific signal (Fig 2D). While smooth in nature, inhibitory predictive remapping is clearly evident in model-internal dynamics and aligns with the ideal inhibition patterns (extracted from ground truth data, Fig 3F/G for a quantification). It should be noted that the predictions here are computed on held out test images, and are thus not based on memorisation of a given scene. Rather, inhibition is dynamically targeted in response to efference copy signals.

**Figure 3:**
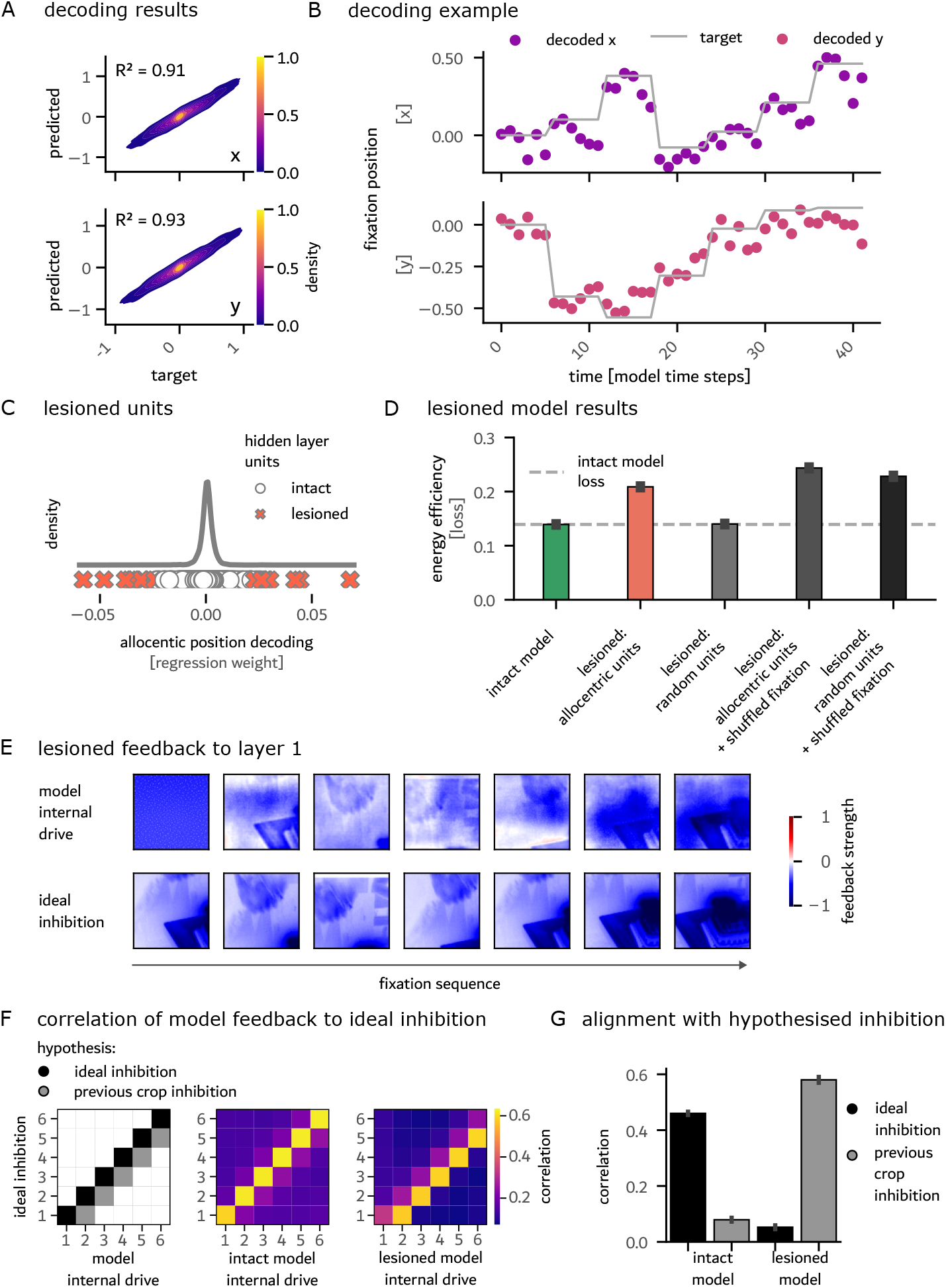
Emergence of an allocentric reference frame that underlies targeted predictive inhibitory remapping. **(A)** Linear decoding of the allocentric fixation X (top) and Y (bottom) coordinates from the RNN hidden layers. The model integrates the relative egocentric saccade commands into an allocentric reference frame. **(B)** Example fixation trajectory (7 fixations with 6 RNN timesteps each) and the decoded coordinates. **(C)** Units targeted during in-silico lesioning were chosen based on their contribution to allocentric coordinate decoding; 0.5% of units with highest betas were selected. **(D)** Lesioning allocentric units led to a significant increase in energy consumption, the extent of which is not matched by random lesioning. **(E)** Example of altered feedback to Layer 1 following the lesioning of allocentric coding units. **(F)** Correlation of the intact and lesioned model feedback with the ideal inhibition targets. **(G)** Correlation of matrices for observed and hypothesised prediction patterns. The internal drive of the lesioned model aligns with the current crop, whereas the intact model aligns with the ideal prediction of the future input.

In conclusion, our RNN outperforms all controls in terms of energy efficiency and exhibits targeted inhibition in a scene-specific manner that is based on efference copy information.

### Allocentric coding emerges from energy efficiency by integrating over sequences of egocentric efference copies

In trying to understand the RNN’s internal mechanism for its inhibitory predictive remapping, we hypothesised that the model may integrate the efferent copies, which are provided relative to the current fixation positions, into an allocentric reference frame that encodes absolute fixation position. To test for this possibility, we trained a linear diagnostic readout to decode the allocentric gaze coordinates from the unit activations in the two hidden RNN layers. This revealed that, indeed, RNN training for energy efficiency led the model to re-code the relative efference copy information into an allocentric code (xcoordinate decoding: *R*^2^ = 0.91, y-coordinate decoding: *R*^2^ = 0.93, see Fig 3A and 3B). Notably, the successful readout relied on only a fraction of all units, as revealed by an analysis of the regression weights (Fig 3C). The high accuracy of the allocentric readout suggests that the model indeed constructed a stable allocentric reference frame based on egocentric efferent copies of previous fixation locations.

### Allocentric reference frame units operate as prediction units

To test for the computational role of the units coding an allocentric reference frame, we performed an in-silico lesioning study on the trained model units. Due to the relatively small number of allocentric-coding units, we only lesioned 0.5% of all units (20 of 4096) by setting them to zero during runtime (see Fig 3C). As a control to this targeted lesioning experiment, we randomly selected the same number of units in the network for lesioning. Random lesioning did not lead to a notable change in loss compared to the intact model. However, lesioning the units that coded for the allocentric position, selected as having large weights in our diagnostic readout, led to a significant increase in the energy efficiency loss (p *<* .001, see Fig 3D). The lesioned model can no longer outperform the control model trained without efference copies (p *>* 0.999). Together, these findings demonstrate the importance of the allocentric-coding units for the model’s ability to perform targeted inhibitory predictive remapping. The computational importance of the allocentric units is further underlined by the allocentric-coding units selected for lesioning having a significantly larger activity on the test-set than the average hidden unit (p *<* .001). This heightened activity despite the training for energy efficiency serves as evidence for the functional importance of these units to minimizing energy expenditure.

To better understand how the in-silico lesions affect network computations, we visualised the model-internal drive (Fig 3E) and compared it to the ground-truth ideal inhibition. We observe that the lesioned model re-used the current crop as inhibitory signal, in contrast to the intact model, which performed a targeted inhibition according to the expected future visual input that results from the saccade to a new location (Fig 2D). To quantify this observation, we compared the model-internal drive of both the intact and lesioned models to the ideal inhibitory signals across all time steps (correlation used as similarity measure). We then compared the resulting similarity matrices to two predictions: one representing a case in which the current crop is used as inhibitory signal, and one where the prediction is aligned with the ideal inhibition (see Fig 3F and Fig 3G). The model-internal inhibitory drive of the intact model aligned well with the ideal future inhibition (*r* = .46, p *<* .001). In contrast, following the targeted lesioning of the allocentric reference frame units, the model fell back to an internal dynamic in which inhibitions were aligned with the current crop (*r* = .58, p *<* .001). Only a minimal alignment with the ideal inhibition remains after the lesioning (*r* = 0.05, p *<* .001). This analysis demonstrates that the intact model relied on the allocentric units to perform targeted inhibition of the expected input that results from the saccade to a new location.

## Discussion

### Energy efficiency and allocentric coding

#### Energy efficiency drives predictive remapping and allocentric coding

We investigated the potential origins of predictive remapping across human-like saccade sequences. Using an RNN model system, we observed that targeted inhibitory predictive remapping emerged naturally from optimizing model computations for energy efficiency. Our diagnostic readout analyses and lesion studies revealed that the model, in an effort to save energy, learned to code in an allocentric reference frame based on the relative efference copies provided to it, and used the corresponding allocentric coding units for targeted inhibition. Only a small number of units were found to be coding for allocentric positioning, consistent with sparse coding principles and with previous results that found few prediction units driving world-model-based inhibition in temporal sequence data (Ali et al., 2022). Our study demonstrates energy efficiency as a driving force in the selforganization of neural networks into distinct functional units with specific computational roles.

#### Allocentric coding as for perceptual stability

The emergence of units capable of translating relative to allocentric coding has significant implications for both predictive remapping and spatial memory encoding. Re-coding spatial information in allocentric terms provides the system with a stable reference frame despite constantly changing fixation locations, enabling object information to be encoded consistently. Such a code may not only underlie our stable perception of the world but may also serve as spatial memory of where things are in our surroundings, enabling the system to use the world itself as an external memory for detailed visual information not maintained in working memory (O’Regan, 1992; Somai et al., 2020).

#### Emergence of complex computation from simple principles

The emergence of allocentric units in an unsupervised manner—as a result of energy-efficient coding—is particularly noteworthy. Previous work studying re-coding from relative to allocentric reference frames in path integration tasks explicitly trained supervised RNNs to perform this mapping (Banino et al., 2018). In such supervised approaches, an open question remained regarding how the necessary training data could be obtained by a biological system. Our work suggests that unsupervised objectives, such as energy efficiency, may be sufficient to yield similar computations without requiring explicit supervision signals. This finding supports the broader principle that complex computational mechanisms can emerge from simple constraints. Rather than requiring genetically hard-coded circuits specifically designed for predictive remapping, our results indicate that fundamental physical constraints—like the brain’s limited energy budget—can drive the emergence of sophisticated computational solutions to complex perceptual problems.

#### Shared coding principles for vision and navigation

Our findings align with observations that spatial navigation and visual exploration may utilize similar cortical networks and coding principles (Nau et al., 2018). According to this view, the hippocampal formation would employ similar mechanisms to code for world-centered visual space as it does for navigable space. Additionally, similar to spatial navigation, the hippocampal formation likely interacts with regions such as the posterior parietal cortex (PPC) and retrosplenial cortex (RSC) to convert egocentric viewing coordinates into allocentric ones (Byrne et al., 2007; Clark et al., 2018). Our computational model provides a mechanistic basis for understanding how these transformations might occur and how they could emerge through energy optimization processes.

### Outlook and future research

#### Integrated active vision and navigation frameworks

Future work could focus on integrating our model into a larger framework of active vision involving working memory and task-guided computations. Such a comprehensive framework would allow us to study further computational benefits of emergent allocentric coding, including targeted information integration, object localization, and more efficient memory encoding. This extended framework could provide insights into the functional separation among multiple brain regions involved in saccadic remapping, including the superior colliculus, frontal eye fields, entorhinal cortex, and various visual cortical areas (Duhamel et al., 1992; Graziano et al., 2002; Leigh & Zee, 2015; Sommer & Wurtz, 2006; Sparks, 2002; Wurtz & Hikosaka, 1986; Nau et al., 2018). Such a more elaborate model architecture might also result in more expressive spatial memory abilities than observed in this simple RNN architecture. A particularly promising direction is exploring the relationship between our findings and grid-coding mechanisms found in both spatial navigation (Doeller et al., 2010; Hafting et al., 2005; Sargolini et al., 2006) and primate active vision (Killian et al., 2012; Nau et al., 2018). Our model could be extended to investigate factors that yield explicit grid-coding schemes supporting allocentric coding in visual attention, potentially bridging the gap between navigation research and visual perception studies.

#### Biological constraints on energy and sampling

Further investigation into more biologically realistic modeling approaches promises to be beneficial. While minimizing mean absolute preactivation is a useful proxy encouraging sparse activations and reducing costly over-inhibition, it remains an imperfect measure, as it does not exhaustively capture the complex metabolic processes underlying neural activity (Ali et al., 2022). We acknowledge that while energy efficiency is conceptually simple, our implementation relies on backpropagation through time (BPTT). This learning rule is arguably of limited biological plausibility (see e.g., Lillicrap & Santoro, 2019). Future work should explore more biologically realistic learning rules that allow to approximate energy minimization in a more direct fashion. Similarly, incorporating retina-like sampling methods (see e.g., Lukanov et al., 2021) would move our model closer to biological reality. This enhancement could provide a more realistic visual periphery to better predict forthcoming visual input and mirror the computational challenge faced by the visual system in translating peripheral neural representations into foveal ones (e.g., by accounting for foveal magnification).

#### Distributed information transfer mechanisms

It will be valuable to study how the implicit world model in our RNN, necessary for allocentric coding and targeted inhibition, is represented in this neuroconnectionist system (Doerig et al., 2023). Understanding how information can be transferred from one topographical location to another—required for presaccadic predictions based on peripheral views—would illuminate the recurrent computations spanning hierarchical levels of abstraction.

#### Neuroscientific implications

The modeling work at hand prompts several testable predictions: (i) A small subset of neurons should be dedicated to allocentric spatial coding, with disproportionate impact on visual stability if selectively manipulated; (ii) Neural mechanisms supporting spatial navigation and visual stability should overlap significantly, predicting that patients with hippocampal-entorhinal-retrosplenial damage would show specific deficits in maintaining visual stability across multiple saccades; and (iii) Disrupting allocentric coding mechanisms should selectively impair predictive remapping while preserving basic visual processing. These predictions offer a framework for empirical validation of our model’s core premise that energy constraints naturally drive the emergence of sophisticated visual stability mechanisms.

### Conclusion

In summary, our findings suggest that striving for energy efficiency as a learning objective can yield complex network behaviors, such as targeted inhibitory predictive remapping and allocentric coding of eye positions. These results contribute to a broader paradigm shift in computational neuroscience by demonstrating how fundamental physical constraints can drive the emergence of sophisticated neural computations without requiring genetically hard-coded architectural designs. Our model thereby provides a potential solution to the hard binding problem through simple physical principles: energy-efficient networks can emergently develop mechanisms that support perceptual stability across saccades.

## Methods

### Conceptual framework

#### Input Drive

The fixed excitatory visual input provided to the model, consisting of fixation crops (greyscaled) and saccade coordinates that the network must process.

#### Preactivation

The sum of inputs to a model unit before the activation function is applied. This serves as a proxy for energy consumption that would occur in biological neural systems.

#### Energy Efficiency Loss

The objective function (mean absolute preactivation across units) used to train the network. Minimizing this encourages sparse neural activity, while the fixed excitatory input prevents network shutdown, forcing the development of targeted inhibitory predictions.

#### Internal Drive

Feedback from higher to lower network layers that counteracts upcoming sensory input, quantified as the model’s learned weights applied to unit activations.

#### Efference Copy

Two-dimensional vector (Δ*x*, Δ*y*) modeling the saccadic movement plan as relative positional change in pixel between fixations, provided to the model before each saccade.

#### Egocentric Coding

Representation of positions relative to current fixation point, implemented in our model as relative coordinate inputs that change with each new fixation.

#### Allocentric Coding

Image-centered (rather than fixation-centered) position representation that emerges in our network, enabling consistent spatial encoding across multiple saccades despite changing viewpoints.

#### Dataset and task design

Naturalistic stimuli were sourced from the MSCOCO dataset (Lin et al., 2015), with human-like fixation sequences generated via the Deepgaze III model (Kü mmerer et al., 2022). Each input sequence included seven fixations. A training set of 48236 images was selected with an additional test set of 2051 images. For each image, 10 different fixation sequences were generated. In each training epoch, one sequence with 7 fixations was randomly chosen. The fixation sequences start in the center of the scene. The original greyscaled scenes have a size of 256 × 256 pixels; the fixation crops were selected to be 128 × 128 pixels.

#### Model design

The model architecture consisted of a fully connected RNN with two hidden layers (2048 units each) and lateral connections. For each time step *t* in the sequence, the input to the network was defined as:

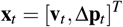

where **v**_*t*_ ∈ **R**^128×128^ represents the flattened visual input (fixation crop) at time *t* and Δ**p**_*t*_ = [Δ*x*_*t*_, Δ*y*_*t*_] ∈ **R**^2^ represents the relative change in fixation position in pixels (efference copy). All components of **v**_*t*_ were strictly non-negative. Analogous to Ali et al. (2022), the input to the model (input drive) was set to be identical to the input image, with no learnable parameters. This ensured that the model could not learn to ignore the input to save energy.

In contrast to Ali et al. (2022), an hierarchical architecture of the model was chosen over a reservoir of units. This change in architecture was done to allow for greater control over the number of parameters and expressivity of the network, as the size of the hidden layers can be easily adjusted to reduce computational complexity in contrast to image-size controlling a large part of the parameter-count through lateral connections. The choice for a hierarchical network also aligns with previous research showing alignment between hierarchical networks and visual cortex (Yamins et al., 2014).

The weight matrices were uniformly initialized following K. He et al. (2015) with weights sampled from 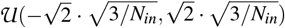, where *N*_*in*_ is the input size of each layer and the scaling factor 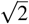 is chosen for ReLU activations. Bottom-up, lateral, and top-down connections operated over six time steps per fixation crop, with the last three steps adding the relative coordinates towards the upcoming fixation location (efference copy). ReLU nonlinearity was applied before computing bottom-up, lateral and top-down connections. Before the first time step of each fixation sequence, a zero-vector was passed to the model to allow for an initial inhibition of the first input.

#### Control models

For the control “no efference copy” (Fig 2A), the network architecture was identical except that the efference copy was not concatenated to the input, reducing the input to **x**_*t*_ = [**v**_*t*_]^*T*^. For the control “smaller crop size” (Fig 2A), the only change to the model design was a reduction of the visual field by 56% to 85 × 85 pixels, resulting in **v**_*t*_ ∈ **R**^85×85^.

#### Training regime

The RNN model was trained to minimise metabolic energy consumption, utilising the mean absolute (L1) preactivation as a loss function. This loss function is formulated as:

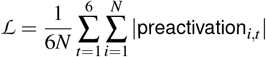

where *ℒ* represents the energy efficiency loss, *N* is the number of units, and |preactivation_*i,t*_ | denotes the absolute preactivation value of unit *i* at timestep *t*. This formulation approximates the biological energy costs associated with neural firing rates and synaptic transmission. The loss was applied across all layers to enforce energy-efficient processing.

Training was done through stochastic gradient descent with the Adam optimiser, a learning rate of 0.0005 and a batch size of 1024. The learning rate was rescaled for each weight matrix by the size of the previous layer as suggested by Roberts & Yaida (2022) to allow for equal contribution of all units to the learning dynamics. This results in an effective learning rate for the bottom-up connections from the first model layer (receiving image and efferent copy data) to the first hidden layer of 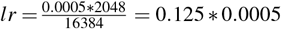 and an unchanged learning rate of 0.0005 for all other weight matrices.

### Analysis

#### Energy-efficiency evaluation

To establish that the learned feedback to layer 1 was inhibitory, we computed the mean feedback drive over all units (128 × 128; see Fig 1B) per timestep and per fixation crop. We then report the average and 99% CI based on the resulting distribution of means. The reported comparisons between model energy efficiency against the control conditions are based on independent ttests that are run over the loss values of all fixations in the test set (N = 2051 scenes x 7 fixations = 14,357). For visualisation (Fig 1A, Fig 2D and Fig 3E), the image with the lowest model loss of all images in the test set that fulfill the criteria of having a standard deviation computed over the pixel values that exceeds 0.25 was chosen. The criteria of a high pixel standard deviation was chosen to allow for an easily visible contrast in the image. The same image and fixation path was chosen for the decoding example (Fig 3B).

To evaluate the effect of the distance of previously visited fixations to the current fixation on the energy efficiency of the network (Fig 2C), we computed the pair-wise distances of all fixations of of all sequences of the test-set and determined the 33.3 and 66.7 percentiles as thresholds for classifying short, medium and long distances between fixations. The losses of the second fixation of all pairs of fixations no more than 3 timesteps apart were classified given the distance and number of time-steps between the fixations. The mean z-scored losses were reported.

#### Targeted in-silico lesioning experiments

To determine the presence of an allocentric coordinate frame, linear decoders were fitted to the 2 normalised hidden layer activations (concatenated over both layers and all model time steps) to predict global fixation coordinates. Subsequently, the impact of specific units on model behaviour was explored through a lesioning analysis. For this, the top 0.05 % of units with the strongest regression coefficients for the global coordinate decoding (total: 20 units) were targeted. As a control, the same amount of randomly selected units was lesioned. To quantify the impact of the targeted lesioning on both the visual precision of the inhibitory feedback and its adaptation to the fixation dynamics, we correlated the observed model-internal drive with the ideal inhibition of the upcoming crop across the full fixation sequence. The ideal inhibition pattern was computed by taking the negative of the exact upcoming fixation patch. The resulting similarity matrix describes the alignment of the model-internal drive at each time point to each of the ideal predictions. Applied to the full test set, we computed 2051 similarity matrices. To see whether the model internal dynamics were related to the saccade targets (i.e. whether the model inhibited the future input), or whether the model used the current fixation patch as inhibition template, we defined two corresponding hypothetical similarity matrices (see Fig 3F). We correlate these two hypothesis matrices with the observed similarity matrices both for the lesioned and intact model. We quantify the fit of the hypothesis matrices by reporting the mean correlation over the test set. Statistical testing was performed against a null-effect with a one-sample ttest (see Fig 3G).

## Acknowledgements

This work was supported by the Deutsche Forschungsgemeinschaft (DFG, German Research Foundation) - 456666331. P.S. and T.C.K. were supported by the ERC Starting Grant TIME (101039524). P.S. was supported by the Federal Ministry of Education and Research (BMBF; Studienstiftung des deutschen Volkes) and the Max Planck Society (MPG; Max-Planck-School of Cognition), and Deutsche Forschungsgemeinschaft (DFG; German Research Foundation) - GRK 2340. The authors thank Dr. Varun Kapoor for assisting with the dataset creation.

## Author contribution

Conceptualization: T.C.K., P.S.; Data curation: T.N.; Formal analysis: T.N.; Funding acquisition: T.C.K.; Investigation: T.N., P.S., T.C.K; Methodology: T.N., P.S., T.C.K.; Project administration: P.S., T.C.K.; Resources: T.C.K.; Software: T.N.; Supervision: P.S., T.C.K.; Validation: T.N., P.S.; Visualization: T.N., P.S., T.C.K.; Writing – original draft: P.S, T.N., T.C.K; Writing – review & editing: P.S., T.C.K., T.N.

## Declaration of LLM usage

Large Language Models (LLMs) were used in limited capacity during the preparation of this manuscript. All scientific content, experimental design, data analysis, interpretation of results, and conclusions are the original work of the authors. The authors take full responsibility for the accuracy and integrity of all content presented in this manuscript. No LLMs were used for data collection or analysis.

